# Neural repetition suppression modulates time perception: Evidence from electrophysiology and pupillometry

**DOI:** 10.1101/2020.07.31.230508

**Authors:** Wouter Kruijne, Christian N. L. Olivers, Hedderik van Rijn

## Abstract

Human time perception is malleable and subject to many biases. For example, it has repeatedly been shown that stimuli that are physically intense or that are unexpected seem to last longer. Two hypotheses have been proposed to account for such biases: one states that these temporal illusions are the result of heightened arousal which speeds up neural clock dynamics, whereas the alternative ‘magnitude coding’ account states that the magnitude of sensory responses causally modulates perceived durations. Common experimental paradigms used to study temporal biases can not dissociate between these accounts, as arousal and sensory magnitude covary and modulate each other. Here, we present two temporal discrimination experiments where flashing stimuli demarcated the start and end of a to-be-timed interval. These stimuli could either be in the same or in a different location, which led to different sensory responses due to neural repetition suppression. Crucially, changes and repetitions were fully predictable, allowing us to explore effects of sensory response magnitude without changes in arousal or surprise. Intervals with changing markers were perceived as lasting longer than those with repeating markers. We measured EEG (Experiment 1) and pupil size (Experiment 2), and found that temporal perception related to changes in event-related potentials (P2) and pupil constriction, both of which have been related to responses in the sensory cortex. Conversely, correlates of surprise and arousal (P3 amplitude and pupil dilation) were unaffected by stimulus repetitions and changes. These results demonstrate that sensory magnitude affects time perception even under constant levels of arousal.

## Introduction

In some of the most critical moments in life, making the right decision hinges on an accurate sense of time: at the start of a runner’s sprint, or in the resolution of a cyclists race, the difference between victory and defeat hinges on choosing the right moment to break away; in our most emotional conversations, the slightest pauses between words can carry a world full of meaning; and during the final resolution of a poker game, the most subtle speedup and slowdown in on opponents’ behavior can be taken as a tell about their odds of having the best hand. But while the clocks on our walls tick away at a steady rate, our ‘internal clocks’ that perceive such intervals are malleable and highly prone to biases, especially under conditions where we are emotional or have a heightened focus and sensitivity.

Identifying the neural mechanisms by which objective time is translated into a subjective experience of time lies at the heart of the study of time perception. To this end, psychophysical studies have attempted to identify the specific conditions where subjective time is systematically over- or underestimated (Eagleman, 2008). These studies generally explain such ‘temporal illusions’ in terms of two possible neural mechanisms. First, converging evidence suggests that the neural ‘clock dynamics’ that give rise to a temporal percept are sped up under heightened states of arousal, which leads to overestimations of perceived durations. In line with this ‘clock-speedup’ view, it has been shown that time is dilated after a train of clicks or flashes (Penton-Voak et al., 1996; Wearden et al., 2009), and that highly emotional stimuli are often perceived as lasting longer (Droit-Volet & Meck, 2007). Further evidence supporting such a mechanism comes from neurophysiological work, which points to striatal dopamine release as a critical modulator of arousal-modulated speedup of clock dynamics (Meck, 1986, 2006; Mikhael & Gershman, 2019; Soares et al., 2016). Second, it has been suggested that the subjective duration of a stimulus is directly affected by the magnitude of the sensory response that it evokes (Eagleman & Pariyadath, 2009; Matthews et al., 2014). This ‘magnitude coding’ account, sometimes referred to as a ‘coding efficiency’ account, is largely motivated by a body of work that demonstrates temporal dilations for stimuli that are known to evoke large responses in sensory neurons. For example, stimuli that are bright, large, and loud are typically perceived as longer than dimmed, soft, and small stimuli (Ekman et al., 1969; Goldstone et al., 1978; Matthews, 2011b; Xuan et al., 2007; Zelkind, 1973). Johnston et al. (2006) was among the first to suggest that magnitude-mediated temporal illusions can be experimentally induced. They showed that a visual stimulus was perceived to last shorter when it was preceded by a flickering adapter stimulus at the same location. As the effects of the adapter were spatially localized, the results favored an explanation based on sensory adaptation over any more global arousal-based mechanism (see Buhusi & Meck, 2009; Mattes & Ulrich, 1998; Shima et al., 2018, who discuss analogous effects of pre-stimulus spatial cueing).

While both mechanisms can account for a range of findings, it has proven difficult to disentangle the two and determine which is the primary driving force between a given temporal illusion. The main reason for this is that the two mechanisms are closely intertwined: fluctuations in arousal and differences in sensory responses strongly covary and affect each other in several ways. Influential theories of perception posit that sensory responses are optimized to signal those stimuli that can be seen as alerting or arousing, either because they violate expectations (Clark, 2013; Friston, 2005; Horvitz, 2000; Rao & Ballard, 1999) or because they are associated with positive or negative reinforcement (Anderson et al., 2011; Cho et al., 2017; Failing & Theeuwes, 2018; Keil & Ihssen, 2004). Conversely, heightened states of arousal can causally modulate the responses of sensory neurons (Buia & Tiesinga, 2006; Mather et al., 2016; Mather & Sutherland, 2011). The intertwined nature of sensory magnitude effects on time perception on the one hand, and alerting- or arousal-based speedup of clock dynamics on the other hand is perhaps best illustrated by research on the temporal oddball paradigm (see Ulrich & Bausenhart, 2019, for a review). Generally, in this paradigm, a sequence of trials is presented in which participants are to estimate the duration of frequent, repeating, standard stimuli and occasional, infrequent oddball stimuli. The typical finding is that the duration of standards is underestimated whereas oddballs appear to last longer. In early studies on this phenomenon, it was interpreted to reflect sped-up clock dynamics, due to the unexpected oddball causing an overall alerting and arousing response (New & Scholl, 2009; Tse et al., 2004; Ulrich et al., 2006). Others, however, have interpreted the temporal oddball effect as a manifestation of magnitude coding (Eagleman & Pariyadath, 2009; Matthews & Gheorghiu, 2016; Matthews et al., 2014; Pariyadath & Eagleman, 2008, 2012). Based on predictive coding theories of sensory processing (Clark, 2013; Friston, 2005; Rao & Ballard, 1999), the repeated standard stimuli were postulated to have an attenuated sensory response, either due to repetition suppression or expectation suppression. The attenuated response would then, in turn, lead to a shorter percept.

Many studies since have focused on disentangling the relative contribution of repetitions and expectations to temporal oddball effects. By presenting stimuli in pairs rather than streams, it was shown that immediate stimulus repetitions themselves also affect perceived durations without a standard/oddball relation (Matthews et al., 2011). By modulating the predictability of stimulus sequences, Cai et al. (2015) found results suggesting that repetitions alone, and not expectations, might be the driving force behind the oddball effect. However, follow-up work demonstrated that the rate of stimulus repetition (Matthews, 2015; Skylark & Gheorghiu, 2017), and the expected sequential position of an oddball both modulate their effect on subjective duration (Wehrman et al., 2018). In addition, it has been shown that the degree to which the oddball physically differs from the standards can scale the temporal oddball effect (Kim & McAuley, 2013; Schindel et al., 2011). Taken together, results suggest that higher-order expectations have the ability to strongly modulate first-order repetition effects. However, they do not directly address the question whether the magnitude of sensory responses is directly responsible for these modifications of subjective time. These findings still leave the possibility that sensory responses either covary with a global sense of contextual surprise, or that they trigger a state of overall alertness or arousal, which in turn speeds up the neural dynamics that are responsible for the sense of time.

Most neurophysiological studies have similarly remained inconclusive as to whether sensory magnitude can directly affect time perception. For example, the amount of temporal dilation in humans in response to different oddballs has been shown to correlate with sensory responses to the same stimuli in the macaque cortex (Sadeghi et al., 2011). Although this would align with the magnitude coding view, this cross-species brain-behavior correlate does not necessitate a causal link between sensory responses and the dilation effect. Noguchi and Kakigi (2006) used magneto-encephalography (MEG) in a design where participants compared the duration of two stimuli that either repeated or changed (cf. Matthews, 2015; Matthews et al., 2011). Switches were perceived as longer than repeats, a dilation effect that correlated across participants with increases in sensory responses. However, a similar relation was found after stimulus onset in climbing neural activity localized to the supplementary motor area, which was interpreted to reflect speed-up in neural clock dynamics. Recently, Ernst et al. (2017) used electro-encephalography (EEG) to show that inter-trial variability in P3 responses predicted the magnitude of the temporal dilation effect. This component was related to the level of surprise or arousal evoked by the oddball (Mars et al., 2008).The authors offered a careful interpretation of their findings, noting that they would fit with both a magnitude coding account as well as a more global arousal-based modulation of clock dynamics.

To our knowledge, only one study has offered neurophysiological support for sensory magnitude effects without manipulating expectation and arousal (Mayo & Sommer, 2013). In this study, macaques judged the interval between two briefly presented visual stimuli as ‘short’ or ‘long’, while local field potentials were recorded from the frontal eye fields (FEF). Neural responses on correctly judged trials were seemingly identical to incorrect trials, except that the sensory responses evoked by the end-marker were weaker or stronger with under- and overestimated durations. As this relation was found at the very end of the interval, it seems unlikely that it can be ascribed to differences in clock speed throughout the interval, and thereby points to a direct interaction between the sensory response and the subjective duration. Nevertheless, as there was no experimental manipulation of sensory responses, it remains unclear what exactly caused these trial-by-trial fluctuations in sensory responses, and thereby what precisely caused the differences in subjective time.

Here, we present results from two experiments that offer new neurophysiological evidence that sensory response magnitude can directly modulate time perception. In both experiments, participants were to time the interval between two briefly presented visual stimuli (cf. Matthews & Meck, 2016; Mayo & Sommer, 2013). These stimuli could be either presented in the same hemifield or in opposite hemifields, but crucially, the location of the end marker was always fully predictable. Nevertheless, results indicated that intervals were perceived as longer when the location changed, compared to when it repeated. Our results strongly indicated that this ‘change bias’ reflects how sensory responses were modulated by repetition suppression, which causally affected perceived durations. In Experiment 1, we recorded EEG and analyzed three components in the event-related potential (ERP) that have been associated with clock speeds (CNV), sensory magnitude (P2) and with either arousal or violation of expectations (P3). Results showed that of these, only P2 amplitude was affected by repetitions, and could be used to predict responses on a trial-by-trial basis. In Experiment 2, we replicated the change bias in a simpler experimental setup, and additionally tracked eye movements to refute alternative explanations of the effect. Furthermore, we analyzed pupil sizes in response to repetitions and changes, which lent further indications that the bias was directly related to the magnitude of sensory responses.

## Experiment 1

In Experiment 1, participants compared two intervals spaced by a memory delay. This experiment was originally designed in order to investigate whether maintenance and retrieval of durations from working memory would yield lateralized signals in EEG if the interval was presented on either side of the display. The analyses addressing this question are presented and discussed in a separate article (Kruijne et al., 2021), so as not to dilute the focus of the research question in the present article. Here, we will focus our analysis on one particular aspect of the task: stimulus locations either changed or repeated in a fully predictable manner, but nevertheless had a robust effect on perceived durations.

As noted above, analyses focused on three relevant ERP-components in particular that are found at central electrode sites and have previously been related to time perception, sensory processing, expectations, and arousal. First, we focus on the CNV, a slow-wave negative deflection that develops during an interval. In early work, the CNV was assumed to directly reflect internal clock computations (Macar & Vidal, 2003; Macar et al., 1999). More recently, a more nuanced interpretation has been put forward that the component reflects time-informed preparation for upcoming events and actions (Amit et al., 2019; Boehm et al., 2014; Kononowicz & van Rijn, 2014; Pfeuty et al., 2005; van Rijn et al., 2011). The second component was the central P2, a positive peak around 200–250ms post-stimulus that is associated with pre-attentive sensory stimulus processing (Van der Molen et al., 2012). The P2 amplitude is modulated by physical stimulus properties, such as its brightness (Crowley & Colrain, 2004; Kaskey et al., 1980), and previous studies have suggested a link between the P2 at stimulus offset and perceived duration (Kononowicz & van Rijn, 2014; Tarantino et al., 2010). Third, we assessed the amplitude of the P3, an ERP complex that has been linked to a range of cognitive functions that include temporal decision making (Lindbergh & Kieffaber, 2013), but also more broadly decision making and outcome evaluation (Jepma et al., 2016; Kelly & O’Connell, 2013; Nieuwenhuis et al., 2005). One canonical observation regarding P3 amplitude is that it relates to the degree of ‘surprise’ in response to a stimulus (Mars et al., 2008), and in turn has been related to predict temporal oddball effects (Ernst et al., 2017).

### Method

#### Participants

Data were collected from 24 healthy participants who were compensated either by means of course credits or a monetary compensation of €10/hour. They were recruited through the research participants pool of the faculty of behavior and movement sciences at the Vrije Universiteit Amsterdam.Four participants were discarded on the basis of noisy EEG data, which was characterized by the preprocessing analyses described below: two participants had a too high number of noisy channels (8 and 10 EEG-electrodes out of 64) and two had a too high percentage of discarded data epochs (38% and 52%). None of the remaining participants were excluded on the basis of behavior. To initially assess behavior, we fit a psychometric curve for each participant by means of logistic regression that predicted the proportion of trials where the second interval was judged ‘longer’ than the first interval, as a function of the absolute difference in time between them. The slope coefficients of this regression were within 2SD from the group mean for each of the 20 participants. The final sample contained 20 participants (10 female, ages 19–26, Mean age 21.9).

#### Stimuli and procedure

Participants were seated in a sound-attenuated, dimly lit room at 75cm viewing distance from a monitor (22 inch, Samsung Syncmaster 2233) with 1680 × 1050 resolution and 120 Hz refresh rate. The experiment was programmed and presented using OpenSesame (Mathôt et al., 2012), with the PsychoPy back-end (Peirce, 2007). Figure 1A depicts the stimulus sequence on an example trial, with black and gray colors inverted for visibility. Trials started with a gray fixation cross (0.2°) presented on a black background for 1000ms, followed by the onset of three, horizontally aligned gray placeholder circles (radius 1.97°, one in the center and two at 9.83°eccentricity) around a central gray fixation dot (0.2°). The placeholders and fixation dot stayed on screen until the end of the trial.

**Figure 1.**
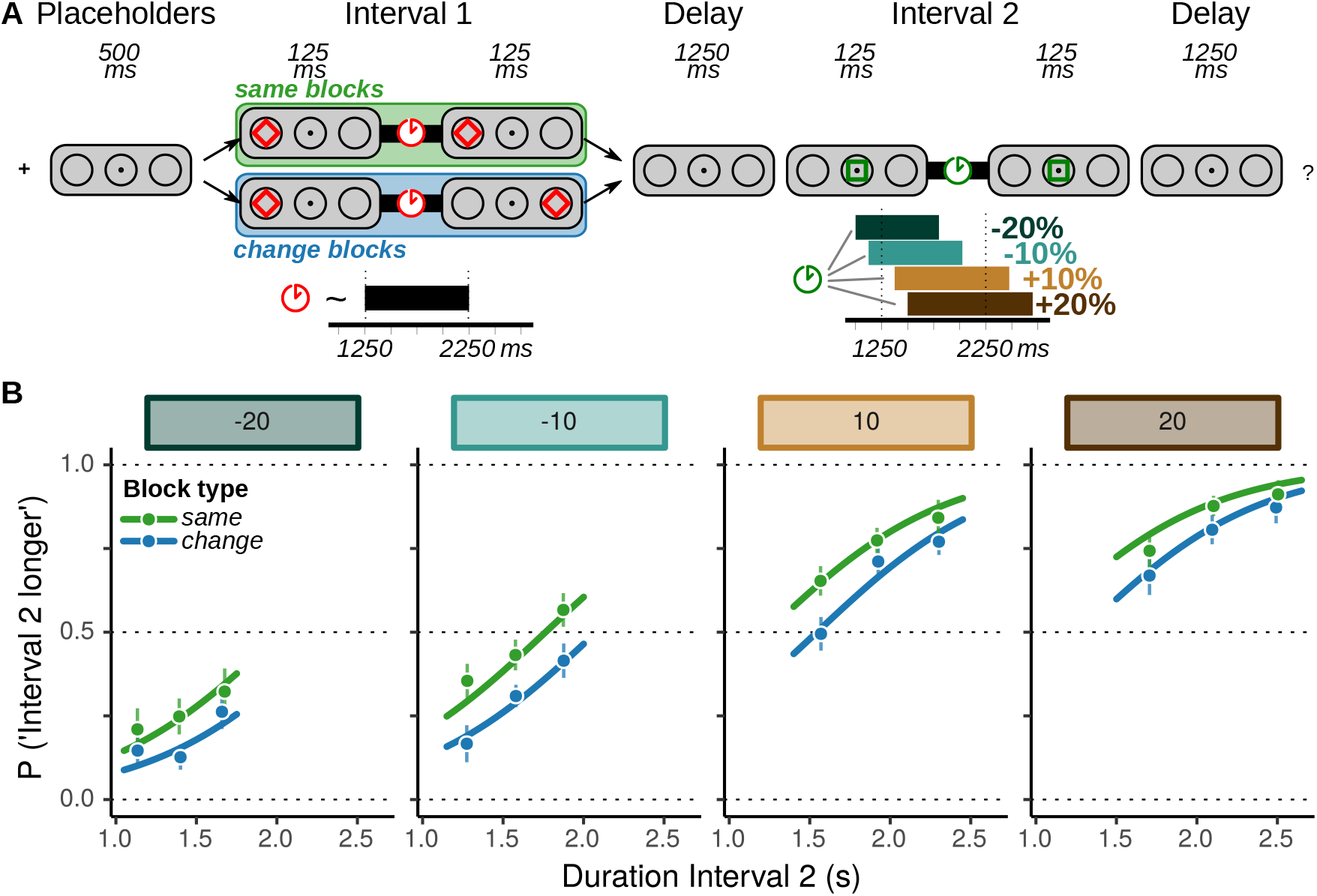
**A** Trial sequence. Participants compared Interval 1, which had a randomly sampled duration, to Interval 2, which was 10 or 20% shorter or longer. Intervals were indicated by flashing markers. Markers for Interval 1 appeared either in the same position (green shading) or changed in position (blue shading), which was varied across blocks. **B** Behavior. Participants’ responses were driven by the relative duration of the intervals, but also by the absolute duration of Interval 2 and by the way Interval 1 was presented. Data points reflect average response rates in per-condition bins of Interval 2 duration, with 95% within subject Confidence interval (Cousineau, 2005; Morey, 2008). Curves reflect the fit of the best GLMM.

The first interval (Interval 1) started 500ms after the onset of the placeholders, which was indicated by a red diamond (3.94°width and height) flashing for 125ms in the left or right circle. The duration of this interval was randomly sampled from a uniform distribution between 1250 and 2250ms. The end of the interval was again marked by a flashing red diamond either at the same or opposite location, dependent on block type (‘same’ or ‘change’ block). Then, a memory delay followed, which lasted 1250ms. After this delay, Interval 2 was presented by means of two green squares flashing in the center circle, one to mark the start and one to mark the end. The duration of this interval was determined relative to the duration of Interval 1, and was always either 10% or 20% shorter or longer. The resulting empirical distributions closely matched the theoretical uniform distributions depicted in Figure 1A (see online Supplemental Information Figure S1 https://osf.io/7gpka/). The offset of Interval 2 was followed by another 1250ms response delay, after which a response screen appeared where they were asked to indicate whether Interval 2 had been shorter or longer by pressing Z or M on a standard keyboard. On each trial, the meaning of these two keys could swap unpredictably, in order to prevent (lateralized) response preparation signals in the EEG. Participants were instructed to be accurate, and that there was no need for a speeded response.

The experiment consisted of eight blocks of 40 trials each. In half of these blocks (‘same’ blocks) the start- and end-marker of Interval were on the same side of the screen; in the other half (‘change’ blocks), they were on opposite sides. Block types alternated throughout the experiment, with an order counterbalanced across participants. Before the start of each block, participants were informed regarding the upcoming block type. Therefore, regardless of block type, participants could never predict the location of the start-marker of Interval 1, but the end marker location was always fully predictable once the start marker had been presented. For Interval 2, both markers were presented in the center of the screen. Participants completed ten ‘same’ and ‘change’ trials as practice’ before the start of the experiment, which were not considered in any of the analyses.

#### EEG acquisition and data cleaning

EEG data were recorded at 512Hz using a BioSemi ActiveTwo system (biosemi.com, Amsterdam, The Netherlands) with 64 channels placed according to the standard 10–20 system. Six additional external electrodes were placed: two were placed on the earlobes to be used as offline reference; two were placed 1cm lateral to the external canthi, and two electrodes 2cm above and below the right eye. From these latter four electrodes, horizontal and vertical electro-oculogram traces (H/VEOG) were constructed by subtracting data from the opposing pairs of electrodes.

Data were preprocessed and analyzed offline using MNE-python (Gramfort et al., 2013; Gramfort et al., 2014) and statistical functions in R. Data were first high-pass filtered at 0.1Hz, removing slow drifts. In order to analyze data epochs with minimal artefacts, we used various algorithms to mark contaminated data for subsequent removal, which are described below. Many of these methods take epochs of equal length as their input, assumed to reflect on-task data of individual trials. To this end we created ‘preprocessing epochs’ with data from −2500ms to 2250ms around the onset of Interval 2. That way, they included a maximum amount of data on-task for each trial, based on the minimum duration of Interval 1, the duration of the memory delay, the minimum duration of Interval 2, and the response delay.

In order to detect channels with noisy data throughout the recording session, we followed the PREP-pipeline (Bigdely-Shamlo et al., 2015) and made use of the RANSAC algorithm implemented in the autoreject package (Jas et al., 2017). This algorithm generates permutations of the dataset where it tries to predict channel data by interpolating the activity from a subset (25%) of channels. If the correlation between the predicted and the observed data is too small (*r* < 0.75) in over 40% of epochs, the channel is marked as faulty. These channels were primarily located at peripheral sites and did not overlap with our electrodes of interest (see below). An overview of marked channels is given in the Supplemental Information, Table S1. These channels were discarded from further preprocessing steps, and are interpolated from surrounding electrodes at the end of data cleaning. Note that this procedure had minimal impact on ERPs, as the analyzed amplitudes were computed from multiple averaged electrodes, and because discarded electrodes had virtually no overlap with our regions of interest.

Data segments contaminated by muscle artefacts were identified by means of a procedure adopted from the PREP-pipeline (Bigdely-Shamlo et al., 2015) and functions in FieldTrip (Oostenveld et al., 2011). Raw data were band-pass filtered at 110-140Hz and the hilbert envelope of the resulting signal was computed. The result was convolved with a 200ms boxcar averaging window, resulting in a per-channel time course estimate of high-frequency power characteristic for muscle activity. We determined a per-channel robust Z-score of such power based on the median and median absolute deviation computed from data within the preprocessing epochs. Data segments where the cross-channel average Z-score was larger than 5.0 were marked as contaminated and not considered in further analyses.

Artefacts caused by eye blinks were detected and filtered out by means of Independent Component Analysis (ICA), which was computed over the full dataset. Since the spatial ICA filters are sensitive to slow drifts, the raw data were first high-pass filtered at 1Hz, and data were subsampled to 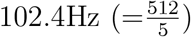 to speed up computation. Independent components were computed using the infomax algorithm (the default in EEGLAB; Delorme & Makeig, 2004). Components were first visually identified as corresponding to blinks, an approach that was subsequently validated by correlating component activity with the VEOG signal during blinks. That is, we filtered the VEOG signal with a 1–10Hz bandpass filter and extracted 1000ms epochs around local maxima. Components that were identified as blink-related, only those components, had a high correlation with VEOG amplitude (*r*^2^ > 0.5).

Horizontal saccades were identified by high-pass filtering the HEOG trace at 1Hz, and convolving the result with a step-wise zero-mean filter kernel of 325ms that went from −1 to 1 by means of a 25ms linear ramp (an approach adopted from ERPlab; Lopez-Calderon & Luck, 2014). In the resulting signal, we identified local maxima that exceeded the 99th percentile as moments of horizontal saccades. Data from the moment of such saccades until the end of a trial were marked as contaminated, and later epochs overlapping with annotated data segments were omitted from analyses.

As a final data cleaning step we applied the autoreject algorithm (Jas et al., 2017) in order to identify unreasonable fluctuations in amplitudes. Autoreject aims to improve on typical epoch rejection based on peak-to-peak amplitudes with a fixed threshold. It uses a bayesian optimization procedure to compute per-channel thresholds, as well as an additional integer value *k.* If a channel’s peak-to-peak value in an epoch exceeds its threshold, autoreject will initially attempt to interpolate its data from neighboring channels. If the number of marked channels in an epoch is larger than *k*, the entire epoch is discarded. An overview of the fitted threshold values is given in the Supplemental Information, together with the number of epochs included after data cleaning (Figures S2 and S3).

#### ERP components: CNV, P2 and P3

Using the algorithms described in the previous section, ‘cleaned’ data epochs were created for ERP analyses. As stated in the introduction, we focused our analyses on data from electrodes at central sites (Cz, C1, C2, FCz, FC1, FC2, CPz, CP1, CP2) where we identified the CNV, the P2 and P3 (see Figure 2A and Figure 3A). In particular, data from these electrodes were spatially averaged and components were defined as the amplitudes in three different time windows in the resulting signal. First, we defined CNV amplitude on each trial as the average amplitude during the 250ms leading up to the end of the interval with respect to a pre-stimulus baseline (for Interval 1, this baseline was 100ms before the onset of the placeholders; for Interval 2 this baseline was 100ms before Interval 2 onset). For the P2, we averaged amplitudes from 200 to 250ms after the offset of each interval. In order to ensure that this measure captured the transient response with minimal interference from the ongoing slow-wave CNV amplitude, the P2 was computed with respect to a 100ms baseline surrounding the moment of interval offset (t=-50ms to +50ms, following Correa & Nobre, 2008; Kononowicz & van Rijn, 2014). The final ERP component of immediate interest, the P3, was defined as the average amplitude between 350 and 400ms using the same baseline as for the P2.

**Figure 2.**
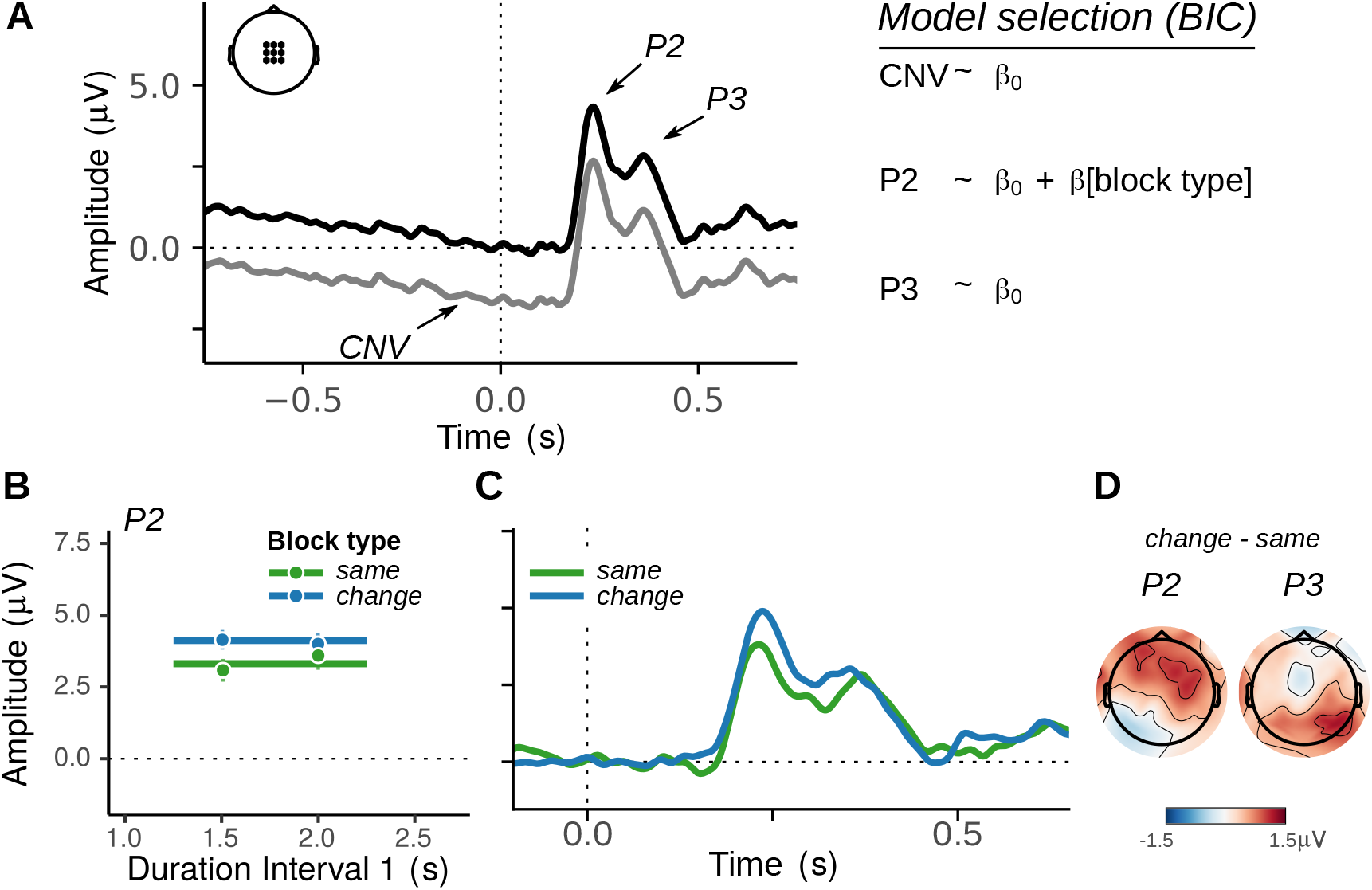
ERPs for Interval 1. **A** Grand-average waveforms at central electrodes. The gray curve has a baseline before the start of the interval, in order to measure the CNV amplitude at the end of the interval. The black curve reflects the same data, but baselined around the presentation of the end marker, to measure the P2 and P3 amplitudes. **B** Model fit, **C** time course, and **D** spatial distribution of the P2 effect. Trials in same- and change blocks evoke P2 components of different amplitudes at central sites, but this difference has disappeared around the time of the P3. Around that time, a contralateral positivity (plotted on the right) was present at occipital sites.

**Figure 3.**
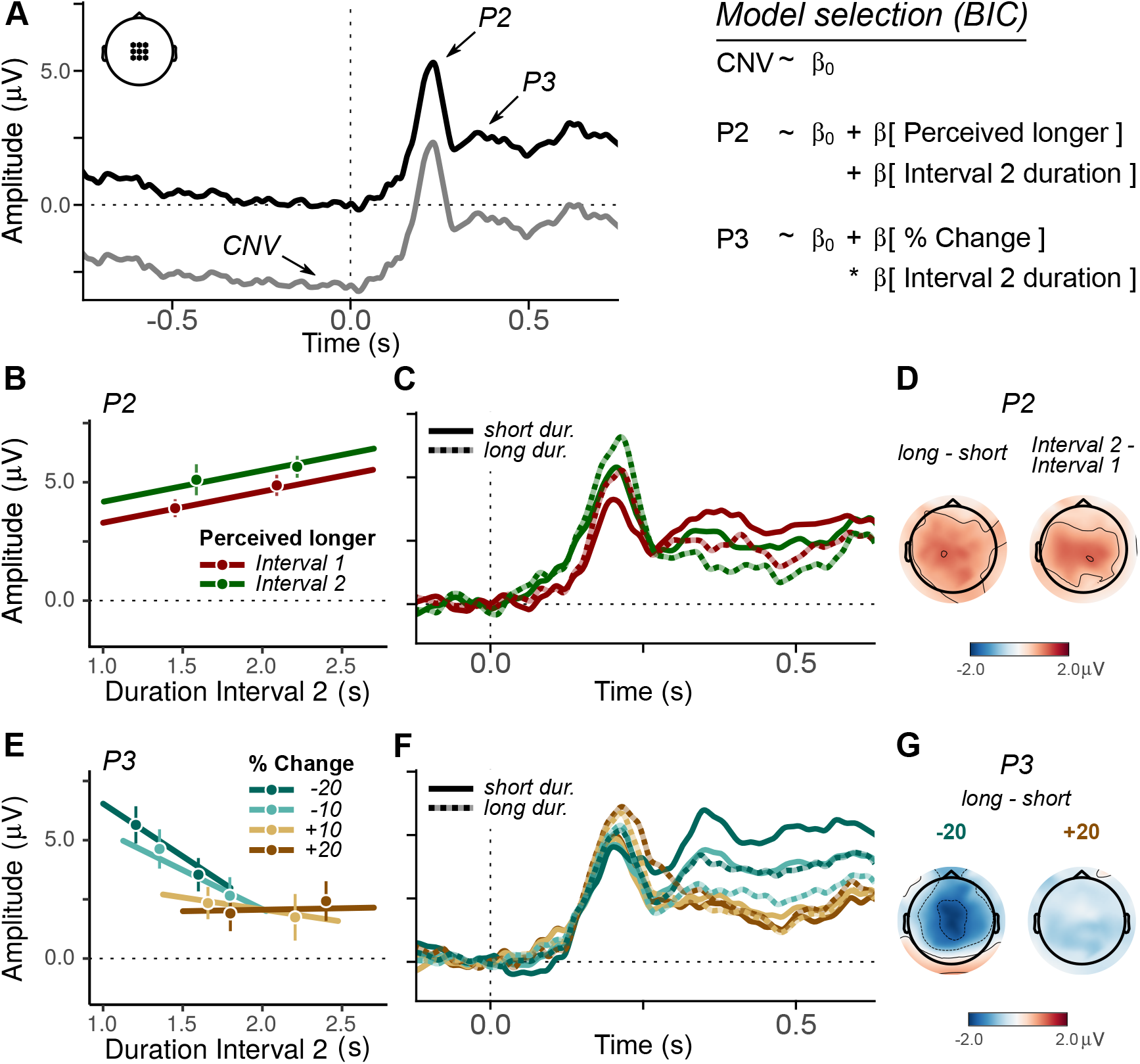
ERPs for Interval 2. **A** Grand-average waveforms at central sites, as in Figure 2, with on the right the best statistical models. P2 amplitudes were predicted by objective duration as well as by the temporal percept. The P3 amplitude was predicted by objective duration interacting with the relative (%) Change **B** Model fits, **C** time course, and **D** Spatial distribution of the P2 effects. Labels ‘long’ and ‘short’ in **C&D** correspond to bins in **B**. **E** Model fit, **F** time course, and **G** spatial distribution of the P3 effects. Again, ‘short’/‘long’ correspond to the bins in **E**

In the main text, we focus on analyses of these components in a data epoch surrounding the end markers, as it is there that we expected to register the effects of the same/change manipulation. Additional analyses specifically addressing neural responses evoked by the start marker are reported in the Supplemental Information (Figure S4 and accompanying text).

#### Statistical analyses

We analyzed behavioral, and electrophysiological data from regions- and time-windows of interest by means of generalized linear mixed models (GLMM) (Baayen et al., 2008). These analyses aimed to characterize the experimental variables that could best account for the behavioral change bias, capture variability in the EEG-data, and identify potential relations between the two. There are several different approaches to statistical inference with GLMMs: for example, one could compare predefined models with prior plausibility on the basis of their fit values, or one might fit a single theory-driven model and determine a p-value for each of its predictors. As the change bias had been an unanticipated finding, we chose a more data-driven model comparison approach. That is, we predefined a set of predictors of interest, and compared the statistical models that could be constructed from these predictors. We base inferences on the BIC, which is typically a more conservative statistic than the AIC or *χ*^2^. We report results regarding the ‘best’ model and its contrast with ‘suboptimal’ models with higher BIC values. In order to additionally report p-values we will report the outcomes of likelihood ratio tests where appropriate.

More precisely, we analyzed behavioral data by means of logistic regression on the response on each trial, and compared models constructed from the following predictors: the duration of Interval 1, the duration of Interval 2, the difference between them (Δ*t*), the percentage change on a trial, and the block type (same/change in marker locations for Interval 1). The most complex model with all predictors and all their interactions was constructed, and as an initial model selection step this model was ‘trimmed down’ using the `lmerTest::step()` function. That is, predictor terms were removed if they did not have additional explanatory value based on AIC-scores. For the resulting ‘trimmed’ model, we fit all less complex nested ‘child’ models, with the only constraint that interaction terms were only included alongside their respective main effects. All models included participant ID as a random intercept term. For each fixed effect in the resulting ‘best’ model, we tested whether including a corresponding random slopes term should be included in the model, and did so if this improved a models’ AIC score.

Each of the ERP components (CNV, P2 and P3 for both Interval 1 and Interval 2) was analyzed using the same model selection procedure, albeit with linear mixed models and with ‘temporal percept’ as an additional predictor (Interval 1 or 2 judged as longer, which was the response variable of the behavioral GLMM). This way, we could establish direct links between ERP-components and behavior. For all analyses, we report the preferred model alongside comparison statistics for suboptimal simpler models where the predictor terms are individually removed We additionally report comparison statistics with other models of theoretical interest. Our conclusions are based on the *ΔBIC*, where we use positive values to indicate that including a term is supported, and negative values for evidence against its inclusion. Wherever possible, we additionally report *p*-values based on the *χ^2^* statistic from likelihood ratio tests.

Note that most factorial analyses, such as traditional ANOVAs or t-tests, would be inadequate for our data, due to the multicollinearity among the predictors of interest. For example, a simple contrast among ERPs for trials where Interval 1 or Interval 2 was perceived as longer would be inappropriate: the latter of these classes would always include more trials where Interval 2 was objectively longer, relatively longer, or where the marker location changed. This similarly impedes most explorations of spatiotemporal differences using cluster-based permutation tests. Comparisons based on the BIC suffer less from this problem as they allow us to assess model complexity as a whole, rather than the estimated contrasts within the model. Nevertheless, this issue of collinearity is to be heeded when interpreting the visualizations and precludes strong conclusions based on resulting model coefficients.

### Results

#### Behavior

Behavior is depicted in Figure 1B, together with the predictions of the model with the best BIC value. This model had three additive fixed effects terms: First, the ‘percentage change’, expressed as a continuous linear term, was a strong predictor of behavior (Δ*BIC* = 15.0; *χ*^2^(1) = 23.8; *p* = 0.001), which describes participants’ ability to correctly perform the task. A random slope was included in the model to capture variability in this term across subjects. Second, participants were more likely to perceive Interval 2 as longer when it was also objectively long (“Duration Interval 2”, expressed as a continuous linear predictor: Δ*BIC* = 296.0; *χ*^2^(1) = 298.8; *p* < 0.001). Note that although participants’ responses would ideally be determined from the relative duration alone, the objective duration is a useful heuristic to determine the correct response which participants might have exploited. Finally, we found that the block type (“same” or “change” block, referring to the markers of Interval 1), affected participants’ responses: Interval 2 was less likely to be perceived as longer than Interval 1 in blocks with a marker change (Δ*BIC* = 82.5; *χ*^2^(1) = 91.2; *p* < 0.001). This effect will hereafter be referred to as the ‘change bias’.

The predictions of this model are depicted in Figure 1B. In order to visualize the fit of binomial response data as a function of a continuous predictor (‘Interval 2 duration’), we split data for each participant across three bins for each condition and computed the average response rate per bin. These averages were used to compute grand averages with 95% confidence intervals of within-subjects effects (Cousineau, 2005; Morey, 2008). Note that this was for visualization purposes only: bins were not used in the analyses.

With respect to our research question it is of particular interest whether the change bias might be accounted for by a sense of ‘surprise’ or violated expectations on change trials. We did not expect this to be the case: the manner of presentation (same/change) was randomly varied across blocks, and therefore participants would have been highly accustomed to repetitions as well as changes. We assessed this in more detail by exploring whether the bias effect was perhaps larger at the start of a block compared to the end. A factorial predictor term ‘block phase’ was defined through a median split on trial index, and was included in a more complex model as a predictor that interacted with ‘block type’. There was, however, no support for this model (Δ*BIC* = –17.5, *χ*^2^(2) = 0.06*, p =* 0.968).

Alternatively, one could hypothesize that even though changes were fully anticipated, these intervals were deemed inherently more interesting than ‘same’-intervals, and thus could have led to an increased state of arousal or alertness throughout their presentation. If this is the case, heightened arousal might have affected neural clock-speed dynamics, which could thereby explain the change bias without a role for sensory repetition suppression. The behavioral results, however, do not support this alternative explanation: if clock-speed were affected by a heightened state of alertness, this would mean that the change bias should be larger for intervals of a longer duration. Such an interaction effect between ‘block type’ and ‘interval duration’, however, had no statistical support (Δ*BIC* = −12.7; *χ*^2^(2) = 4.9; *p* = 0.085). The *χ*^2^ statistic here might marginally hint at a possible interaction, but we note that this was in fact under-additive with a smaller bias for longer durations.

Taken together, the behavioral data support the interpretation that the change bias reflects a constant, additive bias that is driven by differences in the sensory response on ‘same’ and ‘change’ trials. These findings suggest that this bias suggests a sensory magnitude coding effect mediated by repetition suppression, rather than a consequence of sped-up clock computations or an effect mediated by violateds expectations. Next, we investigate whether ERP components lend further support for this view.

#### ERP Components during Interval 1

Figure 2A depicts the overall time course of the selected central electrodes, and highlights the CNV, the P2, and P3 as defined in the methods section. The models that best captured the data are listed in the table on the right. First, for the CNV, we found no evidence that any of the predictors modulated its amplitude: the second-best model included a term for which interval was perceived as longer (Interval 1 or Interval 2), but neither the BIC nor likelihood ratio tests gave an indication that this model was supported (Δ*BIC* = −7.5; *χ*^2^(1) = 0.29). Note that the CNV development at the start of the interval similarly did not differ on same- and change-blocks (Supplemental Information, Figure S4). These results align with the view that neural dynamics underlying subjective time unfold identically on both types of trials, up until the end marker.

For the P2, we did find that it was modulated by the presentation type of the interval. In blocks with a change in marker location, the marker elicited a larger P2 than on blocks with repeating marker locations (Δ*BIC* = 0.15; *χ*^2^(1) = 8.8; *p* = 0.003). This effect is visualized in Figure 2B, accompanied by the ERP time course split for the two block types (Figure 2C). To ensure that this P2 amplitude difference was not partially caused by any potential baselining effects caused by the CNV, we additionally analyzed P2 amplitude in band-pass filtered data (2 to 20Hz), which emphasizes the temporal properties of the P2 while attenuating any modulations by the slower CNV component (following Kononowicz & van Rijn, 2014). Analyses on filtered data similarly supported the model with ‘block type’ as an added predictor (Δ*BIC* = 1.05; *χ*^2^(1) = 9.7; *p* = 0.002). Thus, these results additionally suggest that early sensory responses were larger on trials with a marker change, suggesting that neural response magnitude affected time perception.

In a companion article (Kruijne et al., 2021) we analyzed data from the same experiment and specifically aimed to uncover lateralized signals and their relation to time perception. There, cluster-based permutation analyses indicated that lateralized occipital ERPs were also modulated by block type. We reported a significant difference between 197ms and 455ms post-onset, which we interpreted as a difference in N2pc between these conditions. For the present study, we have reanalyzed the contralateral - ipsilateral difference, averaged in the time window associated with an N2pc (200–300ms). The best LMM was one that included ‘block type’ as a predictor, demonstrating that parieto-occipital, lateralized ERP amplitudes were larger in ‘change’ blocks (Δ*BIC* = 7.9; *χ*^2^(1) = 29.02; *p* < 0.001). In Figure 2D, we plot the average scalp distribution of potentials at 200 to 250ms, and 300 to 350ms, contrasting data from ‘change’ and ‘same’ blocks. These maps are derived from epochs where we mirrored data such that contralateral electrodes are plotted on the right. These maps illustrate how change blocks are characterized by a positivity that peaks more frontally around the time of the P2, but that later in time manifests solely at contralateral, occipital electrodes.

The ERP time course and the scalp distributions depicted in Figure 2C and D suggest that the difference between different trial types is not modulating the central P3 component: around that time the difference is merely found at contralateral occipital electrodes. Indeed, analyses of the P3 found that it was not modulated by any of the predictors, with no support for an effect of trial type (Δ*BIC* = –8.3; *χ*^2^(1) = 0.29; *p* = 0.592) nor for an effect of interval duration (Δ*BIC* = –8.6; *χ*^2^(1) = 0.01; *p* = 0.907). The absence of any modulations of the P3 as reported by Ernst et al. (2017) provides further indication that this change bias is of a different nature than the temporal oddball-effect.

#### ERP Components during Interval 2

In Figure 3A we illustrate the overall time course at central electrode sites around the offset of the second interval. For Interval 2, the CNV was more pronounced, which would align with the view that this component reflects time-based anticipation for the end marker (Kononowicz & van Rijn, 2014; Pfeuty et al., 2005). Nevertheless, we again did not find any support that the CNV amplitude at the end of Interval 2 was modulated by any of the predictors: the best model was the intercept-only model, followed by a model that included the comparison judgment as a predictor (Δ*BIC* = −7.7; *χ^2^*(1) = 0.93; *p* = 0.335). A highly comparable pattern of results was found for the CNV development at the onset of the interval (Supplemental Information, Figure S4).

P2 amplitude for interval 2 was again modulated by experimental variables. That is, the model that best described its amplitude included the predictors ‘Interval 2 duration’ (Δ*BIC* = 2.78; *χ*^2^(1) = 11.4; *p* < 0.001) and ‘perceived longer’ (Δ*BIC* = 0.78; *χ*^2^(1) = 7.9; *p* = 0.005). These predictors were additive, with evidence against an interaction (Δ*BIC* = −8.58; *χ*^2^(1) = 0.06; *p* = 0.798). Again, we reran the analyses on band-passed filtered data to overcome potential carryover effects from the CNV, which resulted in the same statistical model with somewhat stronger evidence (both Δ*BIC >* 1.5; *χ*^2^(1) *>* 10.2; *p* < 0.002). The model fits, the ERP time-course, and topographical distribution of these P2 effects are depicted in Figure 3B-D.

The best model for the P3 was one with the predictors ‘Percentage change’ and ‘Interval 2 duration’, and their interaction (Percentage change: Δ*BIC* = 7.88; *χ*^2^(2) = 25.16; *p* < 0.001, Interval 2 duration: Δ*BIC* = 8.1; *χ*^2^(2) = 25.39; *p* < 0.001, interaction: Δ*BIC* = 7.2; *χ*^2^(1) = 15.8; *p* < 0.001). As illustrated in Figure 3E-G, this interaction entailed that the contrast between ‘short’ and ‘long’ durations of Interval 2 yielded large P3 differences on trials with Interval 2 < Interval 1 (−20% or −10%), but did virtually not differ for trials where Interval 2 > Interval 1 (+10% or +20%).

#### P2 Amplitude predicts the temporal percept

The previous section shows that the P2 at the offset of Interval 2 was larger on trials where Interval 2 was perceived as longer. Out of all explored ERP components discussed above this was the only component with such a relation. To further understand this link, we also assessed whether the inverse held. That is, we assessed whether P2 amplitude was supported as additional predictor in the GLMM predicting behavior, extending the model depicted in Figure 1B. Indeed, we found that the extended model including P2 amplitude as a continuous predictor improved the model (Δ*BIC* = 1.35; *χ*^2^(1) = 9.99; *p* = 0.002). The best model still included the other three predictors (removing either term: Δ*BIC* < –9.78) and interactions between either of the predictors were not supported (Δ*BIC* < –6.87).

The effect of P2 amplitude is depicted in Figure 4, where for each ‘percentage change’ condition the P2 amplitude is divided into three bins. The model prediction with the average P2 amplitude in each bin is plotted as in Figure 1B, but with three levels of transparency reflecting different P2 amplitude bins. The figure illustrates that for certain experimental conditions, participants are approximately 10% more likely to perceive an interval as longer in the highest P2 bin compared to the lowest bin. Note that this figure obscures the observation that generally, the P2 was also found to be larger for longer intervals (Figure 3B).

**Figure 4.**
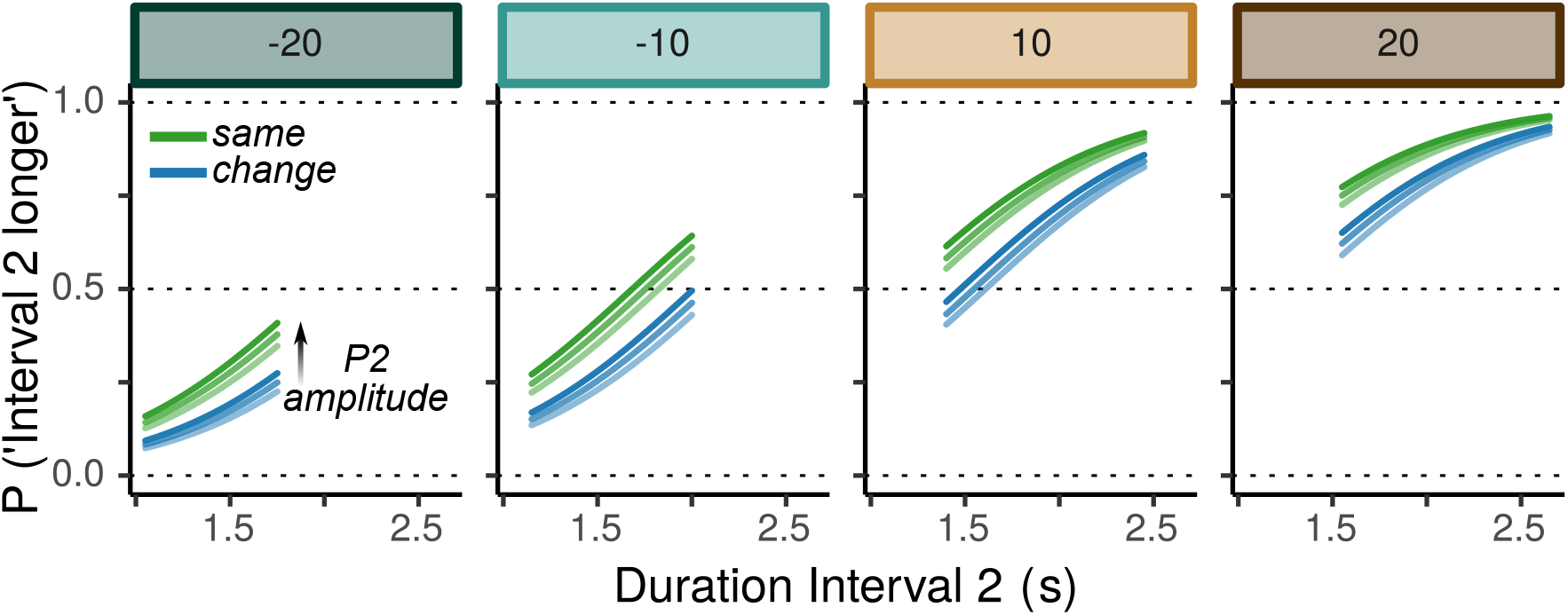
Model predictions for ‘longer’ responses, like the model fit in Figure 1B, but additionally including a predictor for the effect of P2 amplitude. In each condition, the predicted response rates are depicted for three different bins of P2 amplitude. Higher amplitudes are plotted with more opaque lines.

For completeness, we additionally assessed whether the amplitude of any of the other components warranted inclusion in the behavioral model. That is, we tested whether any of the components discussed above would improve the behavioral GLMM. However, none of the components were statistically supported as a predictor for temporal judgments, including the CNV or the P3 at either the onset or offset of Interval 1 and 2.

### Discussion Experiment 1

Experiment 1 revealed that temporal perception was affected by the manner in which Interval 1 was presented: in blocks where the stimuli that marked the start- and end of the interval were in the same location, the time between these markers was perceived as shorter than when the markers were in opposite sides of the visual field. The characteristics of this ‘change bias’ fit with a magnitude coding account of temporal biases, suggesting that neural repetition suppression of the sensory response had a robust, additive effect on perceived duration. ERP analyses established a link between behavior and the P2 amplitude at the offset of both intervals. For Interval 1, blocks with a marker change were associated with a larger P2. For Interval 2, the P2 amplitude on individual trials predicted the upcoming comparison judgement. The CNV and the P3, respectively associated with temporal expectation and surprise, did not differ across trial types, nor did they relate directly to temporal judgments. Results did not support alternative accounts that point to the involvement of either sped-up clock dynamics or any other modulating effects that can be ascribed to differences in higher-order expectations. Therefore, these results offer direct evidence that sensory magnitude, subject to sensory repetition suppression and indexed by the amplitude of the P2, can directly affect the perception of duration.

Even though the results of Experiment 1 establish converging support that the change bias is caused by repetition suppression affecting magnitude coding of temporal durations, several aspects warrant further consideration. First, the experiment was originally not devised to investigate this bias. Rather, the bias was incidentally uncovered uncovered when we analyzed these data to assess representations of duration in working memory. Therefore, our analyses regarding the role of repetition suppression are inherently exploratory and thus requiring subsequent confirmation. Second, it is possible that the requirement to maintain Interval 1 in working memory might have somehow caused the change bias. For example, it might be that the increased sensory response did not affect the *perception* of time, but rather facilitated working memory encoding or retrieval, which subsequently could have led to biased responding in favor of Interval 1. Third, Experiment 1 leaves open the possibility that the bias is not caused by the markers constituting a repetition or change *per se,* but rather somehow is a consequence of the horizontal shift in marker location. For example, the bias might result from inter-hemispheric asymmetries in the processing of time ^1^ (e.g., Bueti et al., 2008; Christman et al., 2003; Harrington et al., 1998; Oliveri et al., 2009; Vicario et al., 2008). Finally, earlier research points to a pronounced effect of saccadic eye movements on duration perception (e.g. Binda et al., 2009; Burr et al., 2010). EOG is potentially not sensitive enough to refute any involvement of eye movements in Experiment 1. Experiment 2 was designed to address these concerns.

## Experiment 2

Experiment 2 was designed to address the concerns outlined above, and thereby offer further confirmatory support that the change bias reflects sensory magnitude coding modulated by repetition suppression. The critical manipulation in Experiment 2 remained the same as in Experiment 1: intervals were presented by means of markers that could either repeat or change. However, the procedure on individual trials was simplified to a bisection paradigm (Allan & Gibbon, 1991; Ulrich & Vorberg, 2009; Wearden, 1991), where participants were to judge a single duration as shorter- or longer than a fixed reference duration, rather than first storing it in working memory for later comparisons. The reference duration in such tasks is assumed to be maintained in long-term memory, but might be sensitive to the history of durations experienced on previous trials (De Jong et al., 2020; Rodríguez-Gironés & Kacelnik, 2001; Taatgen & van Rijn, 2011; Wearden & Bray, 2001). Therefore, change- and same-trials were intermixed within blocks in Experiment 2. Cues at the start of each trial informed participants about the upcoming trial type, and ensured that the location of repeating or changing end-markers remained fully predictable. To test whether the change bias was perhaps limited to horizontal changes we manipulated, across blocks, whether markers would be presented along the horizontal or vertical axis. Finally, Experiment 2 used eye tracking for stringent control for any potential eye movements, both for offline trial rejection as well as online feedback to participants.

We did not record EEG for Experiment 2. Nevertheless, we explored the use of pupil responses as an additional neurophysiological measure to determine the role of sensory magnitude, surprise and arousal in accounting for change bias effects. This is motivated by work establishing that larger pupil dilation is a signature of arousal or surprise (Braem et al., 2015; Satterthwaite et al., 2007). Previous work in both humans and macaques has shown a relation between pupil dilation and temporal behavior (Akdoğan et al., 2016; Suzuki et al., 2016). Conversely, it may be the case that pupil constriction reflexes are informative regarding sensory response magnitude and subject to repetition suppression, although this relation is at this stage only tentative. That is, the pupil constriction response is directly triggered by visual sensory responses (Corneil & Munoz, 2014), and earlier work suggests that constriction responses might be modulated by the magnitude of responses in the sensory cortex (Binda et al., 2013; Mathôt et al., 2014; Naber et al., 2013; Sahraie et al., 2013).

### Method

#### Participants

Data were collected from 32 participants who were recruited through the first-year participant pool of the University of Groningen and had not participated in Experiment 1. The experimental procedure was approved by the Ethical committee of the Faculty of Behavioral and Social Sciences (PSY-1819-S-0019). All participants gave written informed consent before participating, and were treated in accordance with the Helsinki declaration. Data from five participants were omitted due to poor performance (defined below), leaving 27 participants in the final sample (18 female, Mean age 20.6).

#### Stimuli and procedure

Overall trial design of Experiment 2 is depicted in Figure 5A. Participants sat in a dimly lit room in front of a 27” LCD monitor (Iiyama prolite g2773hs-gb1) with a 60Hz refresh rate at 60cm distance from the screen, aided by a chin rest. The experiment started with an instruction phase, where participants familiarized themselves with the reference interval (1750ms) and practiced the task, after which the experimental phase followed.

**Figure 5.**
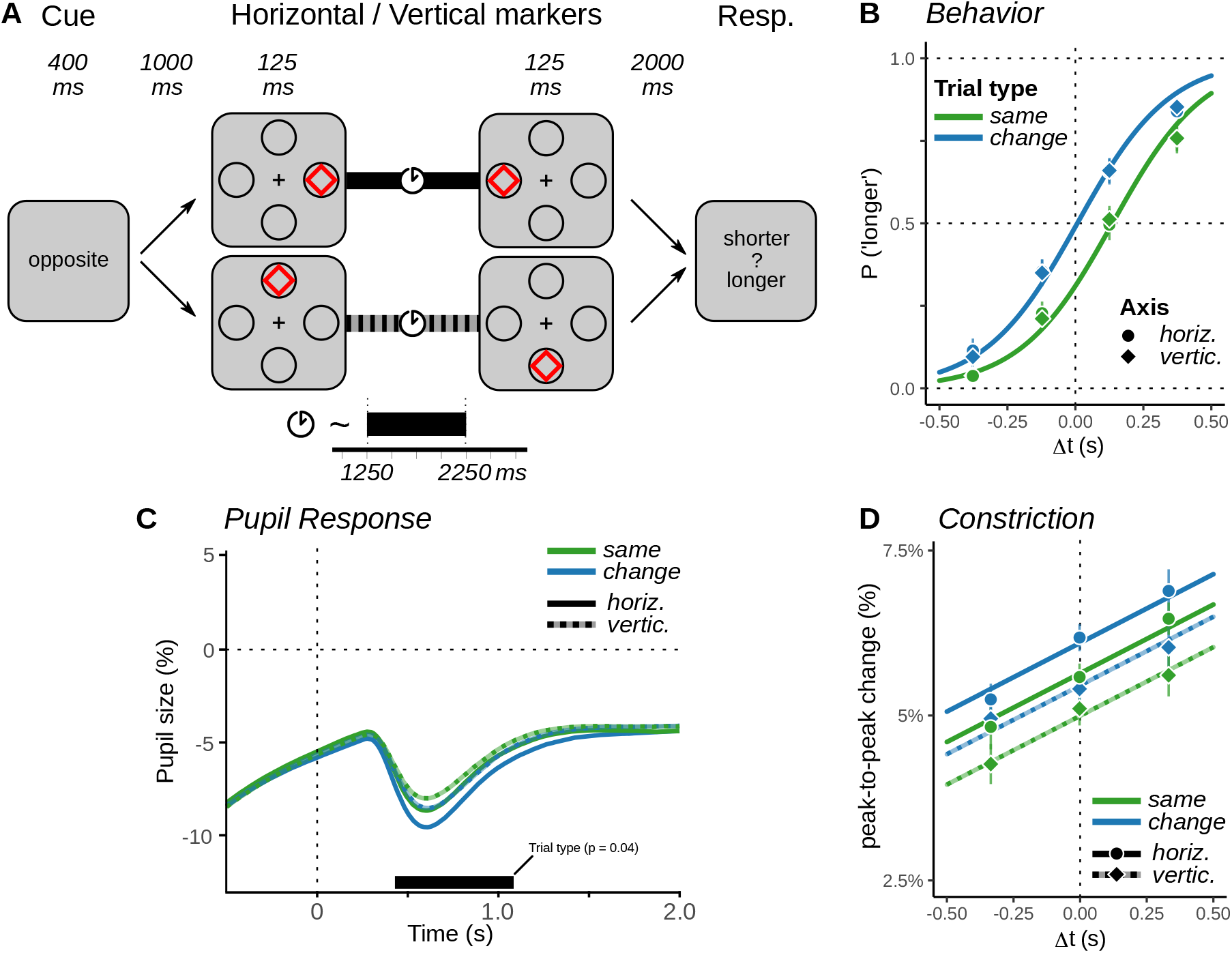
Design and results of Experiment 2. **A** General trial design. Two example ‘change’ trials are illustrated, for horizontal (upper path) and vertical (lower path) blocks. After the cue, the interval was presented through markers on the horizontal or vertical axis. Participants compared a randomly determined interval to the reference duration of 1750ms. **B** Behavior. The x-axis marks deviation (Δ*t)* from the reference interval. As in Experiment 1, ‘change’ trials were perceived as longer. Performance was not affected by the marker presentation axis. Data points and error bars reflect average response in four bins of interval duration, as in Figure 1B. **C** Pupil size in response to the offset of the interval. A significant cluster indicates that the pupil constricts more on ‘change’ trials. **C** The peak-to-peak constriction response at interval offset. Constriction was larger with longer intervals, was larger in horizontal compared to vertical blocks, and larger on change-than on same-trials. Data bins and error bars as in **B**

On-screen stimulus dimensions were identical to Experiment 1, but participants sat 15cm closer to the screen. We report the new, slightly larger visual angles here. Each trial began with a small white fixation cross (0.25°) on a black background. Participants were asked to fixate this cross and press the spacebar, to initiate drift correction and subsequently start a trial. On experimental trials, drift correction was immediately followed by a cue informing participants on the trial type, by means of the word ‘same’ or ‘opposite’, presented for 400ms. In reference trials and practice trials, a 400ms blank screen was shown instead of the cue. This was followed by a 500ms screen with only a fixation dot, after which five white placeholder circles appeared: one in the center, and four at 12.22° eccentricity on the left, right, top, and bottom of the display, each with a 2.45°radius. After another 500ms, the interval was presented as the SOA between the start- and end marker, each flashing for 125ms in one of the placeholders. On each trial, this interval was sampled from a uniform distribution between 1250 and 2250ms. In the experimental phase, markers were red diamonds (4.95°width and height) presented in the peripheral placeholders, either on the horizontal axes or the vertical axes as determined by the block type, and in the same or the opposite location as indicated by the cue. In the instruction phase, start- and end markers were green squares (3.47°) presented within the central placeholder. The placeholders remained on screen for another 2000ms after the end of the interval, after which participants were probed for a response, to indicate whether the presented interval had been shorter (‘F’) or longer (‘J’) than the reference interval (1750ms). They were instructed to be as accurate as possible, without the need to respond fast.

Before the experimental blocks, participants completed an instruction- and practice phase: first, the reference interval was presented nine times to participants, by means of markers that flashed in the center placeholder. Participants were instructed to attend and memorize this duration. Subsequently, they completed ten practice trials with central markers and randomly sampled intervals, where they received trial-wise feedback on their accuracy. The experimental phase consisted of eight blocks of 40 trials each, in which 20 ‘same’ and 20 ‘change’ trials were randomly intermixed. The presentation axis (horizontal or vertical) randomly varied across blocks, with never more than two successive blocks of the same type. Each block started with three ‘refresher’ presentations of the reference interval, after which participants were informed whether markers in the upcoming block would appear on the horizontal or vertical axis in the upcoming block. As in Experiment 1, participants could therefore never predict the exact location of the start marker, but could always infer the location of the offset marker before it was presented.

#### Gaze and pupil data acquisition and analysis

Participants’ right eye was tracked during the experiment using an EyeLink 1000 system (www.sr-research.com), which recorded gaze position in screen coordinates and pupil size (area in arbitrary units) at 1000Hz. Participants were instructed not to blink or move their eyes during a trial, from the cue until the response. If the recorded gaze position was more than 2.45 °from the center and thereby moved outside the central placeholder, participants saw a warning message in orange font at the end of the trial, and the trial was excluded from all further analyses (6.2% of trials).

The analysis of pupil size was conducted offline and followed (Mathôt et al., 2018). First, segments of pupil data recorded between 100 ms before and after a blink were replaced by linearly interpolated data. Two pupil epochs were subsequently created, one around interval onset (−1400 – 1250ms, so as to encompass the time from the start of trial up to the minimal interval duration) and one around interval offset (−500 – 2000ms, encompassing the end of the interval up to the moment of the response). Pupil sizes were baseline-corrected by subtracting the median pupil size in a 200ms window at the start of each trial, and samples were subsequently normalized by dividing pupil size samples by the average median baseline. This resulted in epochs of pupil size expressed as proportional change.

#### Statistical analyses

Overall performance of each participant was assessed by means of a logistic regression that predicted ‘longer’ responses as a function of interval duration, independent of condition. From the regression coefficients, a 50% Just-Noticeable Difference (JND) metric was computed. This JND estimates the duration difference between 25% and 75% ‘longer’ responses, with a larger JND indicating less temporal sensitivity. Participants were excluded if JND*>* 500ms, which would imply that more than half of the trials had fallen within those 25-75% bounds, given the range of sampled durations.

For the remaining participants, behavior was analyzed by means of GLMM’s, predicting responses based on the predictors ‘interval duration’ (numeric predictor), ‘trial type’ (same or change) and ‘axis’ (horizontal or vertical blocks), with a random intercept term and an uncorrelated random slope for interval duration. These two terms described participants’ individual differences in response biases and sensitivity to duration differences, respectively. Based on AIC scores, no other random effects were included. Models constructed from all combinations of predictors and their interactions were again compared using the based on the BIC for statistical inference. For pupil sizes, effects of trial type were initially explored by averaging across trial type (same/change) and presentation axis (horizontal/vertical), and by exploring their average time courses across conditions by means of cluster-based permutation tests. The results, outlined below, suggested that pupil constriction at offset was modulated by trial type. In order to explore this in more detail we additionally computed a per-trial measure of constriction as the peak-to-peak difference in a window 100-700ms after interval offset, which we subjected to model comparisons with LMMs, similar to behavior.

### Results

Data points in Figure 5B illustrate the average proportion of ‘longer’ responses in Experiment 2, computed separately for all levels of ‘trial type’ (same/change) and ‘axis’ (horizontal/vertical), for four bins of interval duration. The curves in this figure reflect the best statistical model based on the BIC. This model had two additive fixed effects: interval duration as a linear predictor (Δ*BIC >* 1000; *χ*^2^(2) = 3163.3; *p* < 0.001), reflective of task performance, and trial type (same/change; Δ*BIC* = 163.2; *χ*^2^(1) = 172.23; *p* < 0.001), which indicated we replicated the change bias effect from Experiment 1. Note that the magnitude of this bias was highly comparable to that reported in Experiment 1, suggesting that the shift to a trial-wise manipulation of marker repetition did not attenuate this bias. Again, we found no evidence for an interaction between duration and trial type (Δ*BIC* = –6.0; *χ*^2^(1) = 2.98; *p* = 0.084). As in Experiment 1, the potential trend hinted at by the *χ*^2^-statistic actually pointed to an under-additive interaction, once again arguing against a clock-speed modulation in change trials. Figure 5B also illustrates that behavior was almost identical in horizontal or vertical blocks. Indeed, there was no support for the additive predictor ‘axis’ (Δ*BIC* = –8.9; *χ*^2^(1) = 0.053; *p* = 0.818) nor for an interaction term that captured a modulation of the change bias by the axis (Δ*BIC* = –17.9; *χ*^2^(2) = 0.061; *p* = 0.970). This illustrates that the change bias was not limited to horizontal displacements, rendering an explanation based on inter-hemispheric communication unlikely^2^.

Pupil sizes at the onset of the interval, did not point to any effects related to the change bias. Exploring the data by means of cluster-based permutation analysis marked one significant cluster that was suggestive of a difference between same- and change trials, from −1150ms to −20ms with respect to interval onset (*p* = 0.002). In this window, change trials were found to have somewhat smaller pupil sizes. Given the timing of this cluster, this likely entails differences in pupil constriction in response to the cue rather than any differences related to temporal perception. Note that change trials were signaled by the word ‘opposite’, and same-trials by the word ‘same’ – the latter of which was smaller and less bright than the former. The permutation test did not identify any significant differences during the interval itself. A LMM predicting the average pupil size in a window of interest at the end of the interval (750 to 1250ms) provided no support for a difference between trial types (Δ*BIC* = –19.5; *χ*^2^(1) = 0.12; *p* = 0.734).

At the offset of the interval, the cluster-based permutation analysis uncovered one significant cluster, for the main effect of trial type, between 429 and 1084ms after interval offset (*p* = 0.044). This was suggestive of more constriction on change trials, as is illustrated in Figure 5C. The traces in this figure also suggest another difference, in that constriction responses were stronger in horizontal blocks compared to vertical blocks. While this difference was indeed reflected in a cluster in the same time range (426 to 771ms), that cluster did not survive permutation correction (*p* = 0.084). We explored this constriction response in more detail with a LMM, but note that it would be misleading to use the pupil size as depicted in Figure 5C as the dependent variable for this LMM. This is because due to the sluggishness of the pupil response, carryover effects from the constriction response to the onset markers can still be observed. Therefore, we instead analyze the constriction response in isolation, by determining the peak-to-peak difference in pupil size in a window 100-700ms following the offset marker.

This peak-to-peak constriction response was best modeled as a function of three additive predictors, as illustrated in Figure 5D. In line with the observations of the cluster test, one of these predictors was the ‘trial type’, with more constriction on change-compared to same-trials (Δ*BIC* = 24.01; *χ*^2^(1) = 45.75; *p* < 0.001). Additionally, model comparisons indicated weaker constriction on ‘vertical’ compared to ‘horizontal’ trials (Δ*BIC* = 67.75; *χ*^2^(1) = 89.5; *p* < 0.001), for which the permutation analysis had also suggested a trend. This aligns with differences in retinal sensitivity along horizontal and vertical axes (Curcio et al., 1990; Seiple et al., 2004). The LMM also allowed us to map the effect of interval duration as a continuous predictor. We found stronger constriction for longer intervals (Δ*BIC* = 284.26; *χ*^2^(1) = 304.96; *p* < 0.001). There was some support for an interaction between ‘axis’ and ‘duration’, though this was not warranted on the basis of the BIC (Δ*BIC* = −13.68; *χ*^2^(1) = 5.56; *p* = 0.018). This interaction (not depicted in Figure 5D) suggested that the duration effect on constriction was somewhat weaker in ‘vertical’ trials compared to horizontal trials. To some extent, the effects of duration might reflect a ceiling-effect on peak-to-peak constriction: on shorter trials, the pupil might already be in a constricted state due to constricting in response to the onset marker. However, this explanation is rendered unlikely given that even for short durations, an effect of trial type is observed, with no support for an under-additive interaction (Δ*BIC* = −17.71; *χ*^2^(1) = 1.53; *p* = 0.216). Instead, the effect of duration on pupil constriction might be similar to what was found for P2 amplitudes in Experiment 1: the effects of constriction might reflect stronger repetition suppression on shorter trials, as well as on ‘same’ trials. Unlike for P2 amplitude, however, we found no support that pupil constriction was predicted by the temporal percept (Δ*BIC* = −21.26; *χ*^2^(1) = 0.01; *p* = 0.905).

### Discussion Experiment 2

Experiment 2 confirmed the behavioral findings of Experiment 1 that a fully predictable stimulus change causes temporal dilation compared to stimulus repetition. The results provided further support for the magnitude coding hypothesis over alternative explanations of the bias, including arousal-based accounts. Behaviorally, the bias was constant across the tested range of durations, which once again suggested that there was no clock speedup in anticipation of an upcoming stimulus change. By strictly controlling for eye movements, both online and offline, we ruled out their potential role in causing the temporal bias. In addition, we found that the bias was virtually identical for horizontal and vertical changes, indicating that this axis does not play a role for the mechanism underlying the bias.

Pupil size measurements further supported a magnitude coding account of the change bias, more so than an account involving fluctuations in arousal. We found stronger pupil constriction responses to the offset marker on change trials. We are not aware of studies that have directly assessed whether neural repetition suppression in the sensory cortex is reflected in the pupil constriction response, and therefore this account remains tentative. Nevertheless, the constriction response has been shown to depend on physical stimulus properties as well as pre-attentive bottom-up salience (Binda et al., 2013; Mathôt et al., 2014). Either way, our findings certainly do not point to an effect of increased pupil dilation, which in previous work has been been associated with motivational salience, arousal, or violation of expectations (Akdoğan et al., 2016; Satterthwaite et al., 2007; Suzuki et al., 2016).

## General Discussion

In two temporal discrimination tasks, we have demonstrated a bias in temporal perception caused by fully predictable changes in sensory input, and have analyzed the behavioral and physiological properties of this bias. We found that a change in the location of a visual start- or end-marker leads to a longer perceived duration of the interval compared to a location repetition. In Experiment 1 we found that this bias in behavior was not modulated by the absolute or relative duration of the intervals, and did not attenuate as participants became more accustomed to changes or repetitions throughout a block: rather, the behavioral bias manifested as an additive effect, consistent across conditions throughout the experiment. ERP analyses indicated that stimulus changes and repetitions were paired with amplified and attenuated central P2 amplitudes, respectively. At the offset of the reference interval, always presented at a central location, the same P2 component was modulated by the interval duration, and was predictive of the upcoming discrimination judgement. These findings strongly relate the P2 amplitude to temporal perception. The P3 measured at the same sites was affected by the objective duration of the reference interval, but was not modulated by the bias nor was it predictive of the upcoming response. In Experiment 2, we replicated the behavioral bias effect in a bisection task, with similar magnitude for horizontal and vertical marker changes. Pupil size analyses showed that the bias was paired with larger constriction in response to change, similar to the P2 modulation in Experiment 1. Taken together, our findings offer direct evidence of magnitude coding effects on time perception. Stimulus repetitions and changes affect the relative magnitude of the sensory response, reflected in the P2 amplitude and pupil constriction, and subsequently affect perceived time. The bias did not seem to be mediated by surprise or arousal, as stimuli were fully predictable and the bias could not be related to P3 amplitude or pupil dilation.

Several differences set our experimental design apart from previous work on temporal biases, in particular from those using oddball paradigms. Most crucially, the sensory changes and repetitions in our task were fully predictable by design, either through blocked presentation or cueing. This allowed us to isolate the effects of repetition independent of surprise. A similar approach has been used previously to dissociate repetition suppression from expectation suppression effects (Larsson & Smith, 2012; Summerfield et al., 2008; Todorovic & Lange, 2012). Another difference is that in our design, stimulus repetitions and changes were physically identical up to the end of the interval, whereas in most oddball designs the deviant stimulus differs from the onset onwards throughout its duration (Matthews, 2011a; Mayo & Sommer, 2013, but see). This means that in oddball designs, sensory magnitude effects could have ongoing effects on time perception throughout the interval, which would give rise to ‘clock speedup’. This would explain why the change bias presented here was characterized by a shift of the psychometric curve, whereas in oddball designs it has often been characterized by a difference in slope(Ernst et al., 2017; Sadeghi et al., 2011; Tse et al., 2004). This interpretation aligns with previous MEG results (Noguchi & Kakigi, 2006) from a study where two intervals were compared that could either be the same or different. Here, it was found that changing stimuli were perceived as longer, and were characterized by a larger visual response at the onset followed by faster changes in climbing neural activity. This climbing neural activity was localized to the supplementary motor area and compared to the CNV-component in EEG recordings.

Regarding the CNV, it may seem remarkable that even though it was clearly present in our task, we did not find that it was related to the temporal bias, nor that it was related to discrimination responses. This extends earlier work that suggests that the CNV does not reflect the timing process itself, but instead marks processes that are contingent on time, such as preparing for upcoming stimuli or actions (Boehm et al., 2014; Kononowicz & van Rijn, 2014; van Rijn et al., 2011). In both experiments, several aspects of our design might have prevented such preparation: Intervals were uniformly distributed across a relatively wide range, participants were probed to respond only late after the offset of the interval, and the mapping of the percept to either of the keys was not known to the participant during estimation. Together, these factors might have prevented preparation-effects on the CNV that previous studies interpreted as a marker of the internal clock (Hiltraut et al., 2003; Macar & Vidal, 2003; Trillenberg et al., 2000).

The P3 on the other hand, did seem to capture aspects of the temporal comparison. At the end of Interval 1, the P3 was constant in amplitude across experimental factors, in line with our assumption that stimulus changes and repetitions were both fully anticipated. However, when the temporal comparison was to be made at the end of Interval 2, P3 amplitude was smaller for longer durations (cf. Lindbergh & Kieffaber, 2013), interacting with duration relative to Interval 1. This aligns with the classical interpretation of the P3 as index of surprise (Mars et al., 2008), but in addition points to the relation between the P3 and decision making processes (Twomey et al., 2015). Taken together, these results suggest that the P2 and the P3 might index different aspects of temporal decision making, which both contribute to the later comparison judgment: whereas the P3 seems primarily driven by the relative duration, thus reflecting the comparison that a participant needs to make, the P2 appeared to index the overall sense of objective duration.

Thus far we have phrased our findings within the context of the magnitude coding account: stimulus repetitions and changes cause attenuation or enhancement of sensory responses due to neural repetition suppression, which causally affects the perception of time. A subtly different interpretation of our findings would be that neural repetition suppression itself reliably depends on the time between repetitions, and that the resulting sensory magnitude is thus used by the brain as a heuristic to tell time. Not many studies have explored the effects of inter-stimulus intervals on repetition suppression, but some findings in the rodent auditory cortex (Budd et al., 2013) and the human visual cortex (Noguchi et al., 2004) suggest that sensory response magnitude indeed monotonically increases as a function of this duration. This implies that the amount of repetition suppression to two subsequent stimuli could in theory be used to infer how much time has elapsed between them. Although our results might fit this interpretation, future research is necessary to fully understand the temporal dynamics of repetition suppression effects and whether they indeed contribute to our perception of time.

By dissociating stimulus repetition from stimulus expectation, we were able to identify a temporal bias that we could attribute to neural repetition suppression. We emphasize, however, that the inverse relation might not always hold, as not every form of repetition will necessarily affect temporal percepts. The precise effects of stimulus repetition are known to vary across stimuli (Amado & Kovács, 2016) and the cortical area under consideration (Larsson & Smith, 2012), to the extent that certain configurations demonstrate repetition enhancement rather than suppression (de Gardelle et al., 2013; Segaert et al., 2013). It is to be expected that not every sensory area plays a crucial part in temporal cognition and in computing perceived durations. Therefore, the change bias paradigm might be used to provide critical insights into which sensory systems do contribute to time perception, as they might isolate sensory repetition suppression effects without confounding these with modulations through arousal or surprise. Previous research suggests that ‘change biases’ similar to the location-effects presented here could arise with changes in size (Matthews, 2011a) or changes in the direction of two arrowheads (Noguchi & Kakigi, 2006). Visual processing of stimulus location, size, and direction is classically attributed to the dorsal ‘where’ pathway (Haxby et al., 1991). These temporal change biases therefore converge with recent findings from neuroimaging that the dorsal pathway, involving the fronto-parietal network in particular, plays a critical role in perceiving the timing of visual events (Battelli et al., 2008; Harvey et al., 2020; Harvey et al., 2015; Walsh, 2003). Future neuroimaging work might help identify the neuroanatomical substrates that underly the modulations in EEG- and pupillometry reported here.

In conclusion, we believe that our findings present compelling evidence that sensory repetition effects affect human time perception, even when repetitions and changes are fully in line with expectations. Results from behavior, electrophysiology and pupillometry all supported this interpretation, demonstrating that neural magnitude can modulate time perception in the absence of arousal-based modulations of neural clock dynamics. In neuroimaging, repetition effects sensory neurons have a strong tradition of being used as a tool to map out neural tuning in cortical areas, uncovering the pathways that give rise to a percept (Grill-Spector et al., 2006). In the same vein, temporal biases like the change bias can be used to determine how low-level sensory responses give rise to a temporal percept.

## Footnotes

1 behavioral data did not immediately support this hypothesis: We found no support for an interaction effect of marker location/direction and block types (Δ*BIC* = −16.1; *χ*^2^(2) = 1.55; *p* = 0.460).

2 Furthermore, just as for Experiment 1 we found no evidence for asymmetrical change biases contingent on marker locations in horizontal blocks (Δ*BIC* = –10.9; *χ*^2^(2) = 5.68; *p* = 0.058).

